# netNMF-sc: Leveraging gene-gene interactions for imputation and dimensionality reduction in single-cell expression analysis

**DOI:** 10.1101/544346

**Authors:** Rebecca Elyanow, Bianca Dumitrascu, Barbara E. Engelhardt, Benjamin J. Raphael

## Abstract

**Motivation:** Single-cell RNA-sequencing (scRNA-seq) enables high throughput measurement of RNA expression in individual cells. Due to technical limitations, scRNA-seq data often contain zero counts for many transcripts in individual cells. These zero counts, or *dropout events*, complicate the analysis of scRNA-seq data using standard analysis methods developed for bulk RNA-seq data. Current scRNA-seq analysis methods typically overcome dropout by combining information across cells, leveraging the observation that cells generally occupy a small number of RNA expression states.

**Results:** We introduce netNMF-sc, an algorithm for scRNA-seq analysis that leverages information across *both* cells and genes. netNMF-sc combines network-regularized non-negative matrix factorization with a procedure for handling zero inflation in transcript count matrices. The matrix factorization results in a low-dimensional representation of the transcript count matrix, which imputes gene abundance for both zero and non-zero entries and can be used to cluster cells. The network regularization leverages prior knowledge of gene-gene interactions, encouraging pairs of genes with known interactions to be close in the low-dimensional representation. We show that netNMF-sc outperforms existing methods on simulated and real scRNA-seq data, with increasing advantage at higher dropout rates (e.g. above 60%). Furthermore, we show that the results from netNMF-sc – including estimation of gene-gene covariance – are robust to choice of network, with more representative networks leading to greater performance gains.

## Introduction

Single-cell RNA-sequencing (scRNA-seq) technologies have provided the ability to measure differences in gene expression between organisms, tissues, and disease states at the resolution of a single cell. These technologies combine high-throughput single-cell isolation techniques with next-generation sequencing, enabling the measurement of gene expression in hundreds to thousands of cells in a single experiment. This capability overcomes the limitations of microarray and RNA-seq technologies, which measure the average expression in a bulk sample, and thus have limited ability to identify differences in gene expression in individual cells or subpopulations of cells present in low proportion in the sample Wang et al. [2009].

The advantages of scRNA-seq are tempered by undersampling of transcript counts in individual cells due to inefficient RNA capture and low numbers of reads per cell. The result is a gene × cell matrix of transcript counts containing many *dropout events* which occur when no reads from a gene are measured in a cell, even though the gene is expressed in the cell. The frequency of dropout events depends on the the sequencing protocol and depth of sequencing. Cell-capture technologies such as Fluidigm C1 sequence hundreds of cells with high coverage (1-2 million reads) per cell, with dropout rates ≈20−40% [Ziegenhain et al., 2017]. Microfluidic scRNA-seq technologies, such as 10X Genomics Chromium platform, Drop-Seq, and inDrops sequence thousands of cells with low coverage (1K-200K reads) per cell, and thus high rates of dropout (up to 90%) [Zilionis et al., 2017]. Moreover, transcripts are not dropped out uniformly at random, but in proportion to their true expression levels in that cell.

In recent years, multiple methods have been introduced to analyze scRNA-seq data in the presence of dropout events. These methods address one or more of the first three steps that constitute most scRNA-seq pipelines: (1) imputation of dropout events; (2) dimensionality reduction to identify lower-dimensional structures that explain most of the variance in the data; (3) clustering to group cells with similar expression. Imputation methods include MAGIC [Van Dijk et al., 2018], a Markov affinity-based graph method, scImpute [Li and Li, 2018], a method that distinguishes dropout events from true zeros using dropout probabilities estimated by a mixture model, and SAVER [Huang et al., 2018], a method that uses gene-gene relationships to infer the expression values for each gene across cells. Dimensionality reduction methods include ZIFA [Pierson and Yau, 2015], a method that uses a zero-inflated factor analysis model, and SIMLR [Wang et al., 2017], a method that uses kernel based similarity learning for dimensionality reduction. Clustering methods include BISCUIT, which uses a Dirichlet process mixture model to perform both imputation and clustering [Azizi et al., 2017], and CIDR, which uses principal coordinate analysis to cluster and impute cells [Lin et al., 2017b]. Other methods, such as Scanorama, attempt to overcome limitations of scRNA-seq by merging data across multiple experiments [Hie et al., 2018]. Supplemental Table S1 gives a list of these and other related methods.

We introduce a new method, netNMF-sc, which leverages prior information in the form of a gene interaction, protein interaction, or gene co-expression network during imputation and dimensionality reduction of scRNA-seq data. netNMF-sc uses network-regularized non-negative matrix factorization (NMF) to factor the transcript count matrix into two low-dimensional matrices: a gene matrix and a cell matrix. The network regularization encourages two genes connected in the network to have a similar representation in the low-dimensional gene matrix, recovering structure that was obscured by dropout in the transcript count matrix. The resulting matrix factors can be used to cluster cells and impute values for dropout events. While netNMF-sc may use any type of network as prior information, a particularly promising approach is to leverage tissue-specific gene co-expression networks derived from the vast repository of RNA-seq and microarray studies of bulk tissue, and recorded in large databases such as Coexpressdb [Okamura et al., 2014], Coexpedia [Yang et al., 2016], GeneSigDB [Culhane et al., 2009], and others [Lee et al., 2004, Wu et al., 2010].

We demonstrate netNMF-sc’s advantages in recovering cell clusters in both simulated and real scRNA-seq datasets with high dropout. We also show that netNMF-sc outperformed several existing methods in recovering differentially expressed marker genes and gene-gene correlations following imputation of dropout events. netNMF-sc provides a flexible and robust approach for incorporating prior information in imputation and dimensionality reduction of scRNA-seq data.

## Results

### netNMF-sc algorithm

Let **X** ∈ ℝ^*m×n*^ be a matrix of transcript counts for *m* transcripts measured in *n* single cells obtained from an scRNA-seq experiment. It has been observed that the majority of variation in transcript counts is explained by a small number of “expression signatures” that represent cell types or cell states. Since **X** is non-negative, one can use non-negative matrix factorization (NMF) [Lee and Seung, 1999] to find a lower dimensional representation by factoring **X** into an *m* × *d* gene matrix **W** and a *d* × *n* cell matrix **H**, where *d* ≪ *m,n*, and the elements of both **W** and **H** are non-negative. We formulate this factorization as a minimization problem,

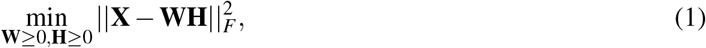

where ||⋅||_*F*_ is the Frobenius norm, and ≥ indicates non-negative matrices whose entries are ≥ 0.

Because of high rates of dropout and other sources of variability in scRNA-seq data, the direct application of NMF to the transcript count matrix **X** may lead to components of **W** and **H** that primarily reflect technical artifacts rather than biological variation in the data. For example, Finak et al. [2015] observe that the number of dropped out transcripts in a cell is the primary source of variation in several scRNA-seq experiments.

To reduce the effect of dropout on the factorization, we propose to combine information across transcripts using prior knowledge in the form of a gene interaction network, and to include a zero inflation term in the model. We incorporate network information using graph regularized NMF [Cai et al., 2008] which includes a regularization term to constrain **W** based on prior knowledge of gene-gene interactions. In addition, we expect that many of the zero entries in **X** do not represent zero levels of that transcript but rather the effect of dropout. We include zero inflation using a binary matrix **M** that masks zero entries in **X**, such that a non-zero entry in *a*_*ij*_ in **WH** is not penalized when the corresponding entry *x*_*ij*_ of **X** is equal to 0 (see Methods for further details). The resulting method, netNMF-sc, performs matrix factorization by solving the following optimization problem:

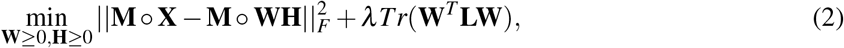

where *λ* is a positive real constant, **L** is the Laplacian matrix of the gene-gene interaction network, ∘ indicates element-wise multiplication (or Schur product of matrices), and *Tr*(⋅) indicates the trace of the matrix.

netNMF-sc uses the resulting matrix **H** to cluster cells, and the product matrix **WH** to impute values for dropout events in the transcript count matrix (Figure 1). We select the regularization parameter *λ* by assessing clustering performance on simulated data (Section S2).

**Figure 1:**
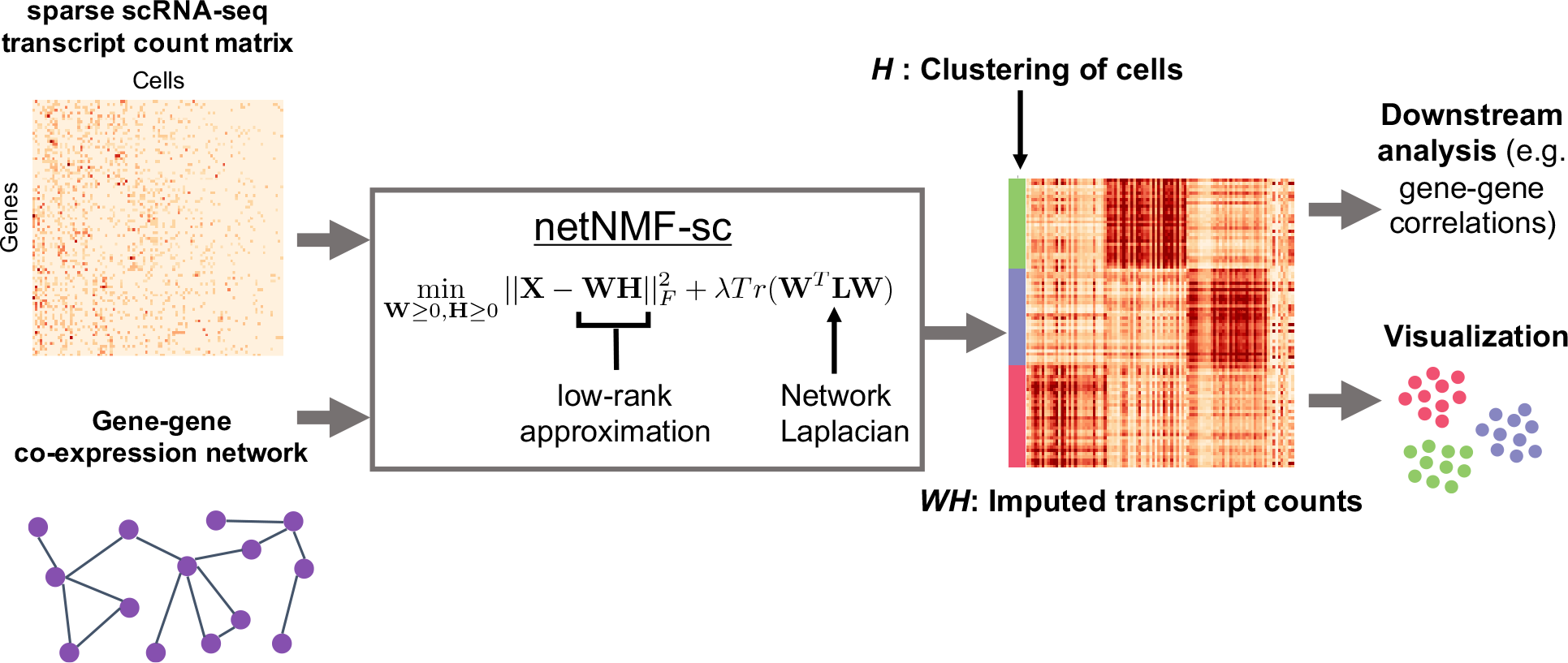
Overview of netNMF-sc. The inputs to netNMF-sc are: a transcript count matrix **X** from scRNA-seq data and a gene network. netNMF-sc factors **X** into two *d*-dimensional matrices, gene matrix **W** and a cell matrix **H**, using the network to constrain the factorization. The product matrix **WH** imputes dropped out values in the transcript count matrix **X** and is useful for downstream analysis such as quantifying gene-gene correlations, while **H** is useful for clustering and visualizing cells in lower-dimensional space.

### Evaluation on simulated data

We compared netNMF-sc and several other methods for scRNA-seq analysis on a simulated dataset containing 5000 genes and 1000 cells and consisting of 6 clusters with 300, 250, 200, 100, 100, and 50 cells per cluster respectively. We generated this data using a modified version of the SPLATTER simulator [Zappia et al., 2017], modeling gene-gene correlations using a gene coexpression network from Yang et al. [2016]. We simulated dropout events using one of two models: a *multinomial dropout model* [Linderman et al., 2018, Zhu et al., 2018] and a *double exponential dropout model* [Li and Li, 2018, Azizi et al., 2017]. Further details are in Methods.

We first compared the clustering performance of CIDR, *k*-means clustering of the (unimputed) count data, and *k*-means clustering of the imputed data from MAGIC, scImpute, NMF, and netNMF-sc. For both CIDR and *k*-means clustering, we set *k* = 6 to match the number of simulated clusters. We ran netNMF-sc with *d* = 10, *λ* = 10 (based on cross-validation) and using the same gene co-expression network [Yang et al., 2016] that was used to create the simulated data. (Note that this does not guarantee that the gene-gene correlations in the simulated data are exactly those in the network, as only a subset of genes are selected and some pairs of differentially expressed genes may not be represented by an edge in the network). For MAGIC, scImpute, and the raw data we performed *k*-means clustering on the first 10 principal components (PCs) of the imputed data in order to reduce the dimensionality to match the dimensionality of the matrix **H** computed by netNMF-sc.

We compared the performance of the methods at dropout rates ranging from 0 (no dropout) to 0.80 (80% of the values in the data are zero). At each level of dropout, 20 simulations were performed. Across all dropout rates, the clusters identified using netNMF-sc had higher overlap with true clusters compared to the clusters identified using other methods (Figure 2(a)). The improvement for netNMF-sc was especially pronounced at higher dropout rates; for example at a dropout rate of 0.7, netNMF-sc achieved an average adjusted rand index (ARI) of 0.81, compared to 0.52 for the next best performing method, NMF (Figure 2(d)). We observe a similar improvement in clustering performance using the *double exponential dropout model* (Figure S6(a)). At a dropout rate of 0.7, netNMF-sc achieved an average ARI of 0.79, compared to 0.41 for the next best performing method, scImpute (Figure S6(d)).

**Figure 2:**
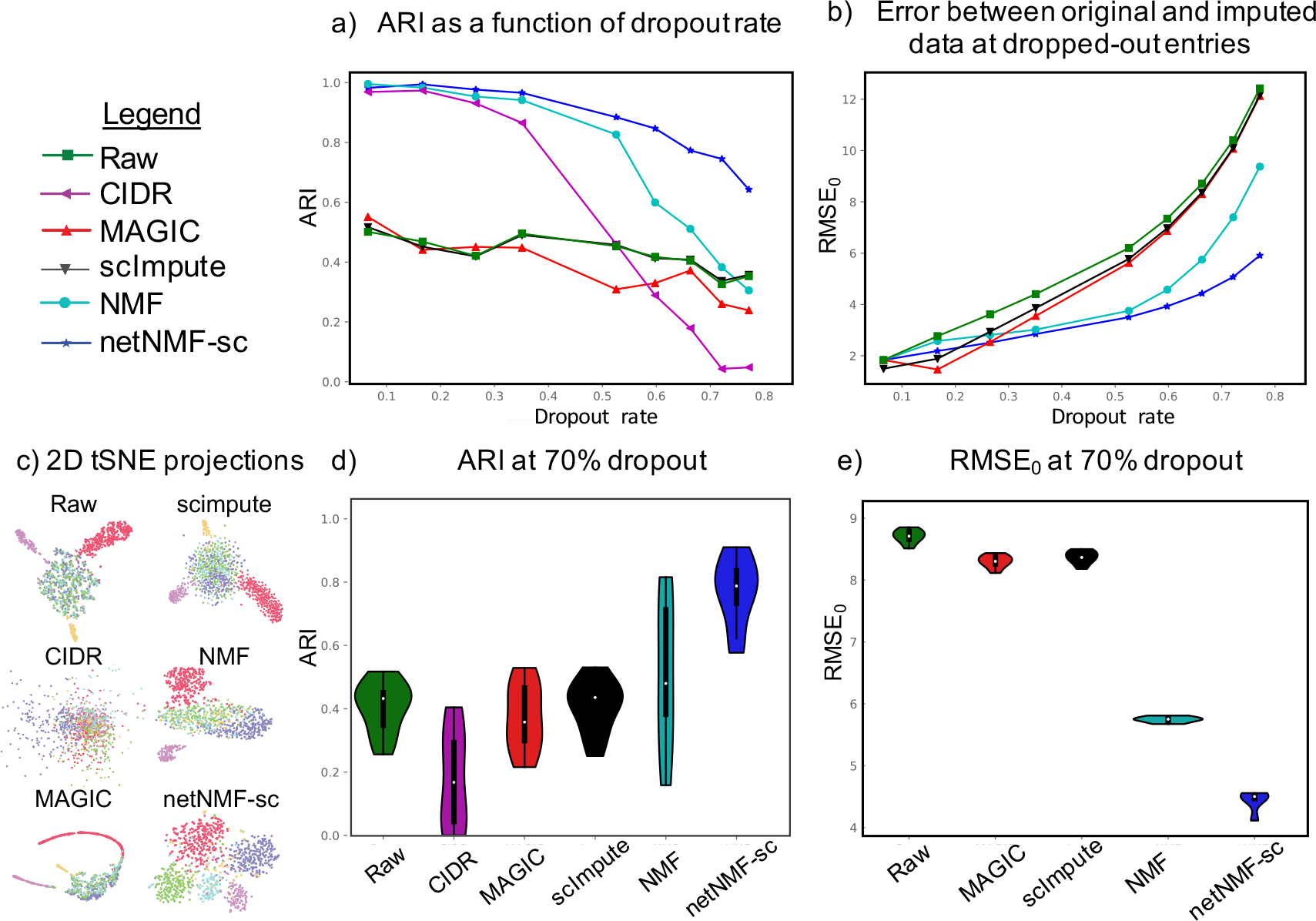
Comparison of netNMF-sc and other methods on clustering and imputation for a simulated scRNA-seq dataset containing 1000 cells and 5000 genes, with dropout simulated using a multinomial dropout model. (a) Clustering results for several scRNA-seq methods on simulated data with different dropout rates. (b) Imputation results with different dropout rates. (c) 2D tSNE projections of imputed data at 70% dropout. (d) Distribution of adjusted rand index (ARI) at 70% dropout over 20 simulations. (e) Distribution of root mean squared error (RMSE_0_) at 70% dropout over 20 simulations.

We compared the performance of netNMF-sc and other approaches on the task of imputation by computing the root mean squared error (RMSE) between **X**′, the simulated transcript count matrix before dropout, and the imputed matrix. We first compute RMSE_0_, the RMSE between **X**′ and the imputed matrix restricted to entries where dropout events were simulated. At low dropout rates (*<* 0.25), netNMF-sc had similar average RMSE_0_ as other methods, but at higher dropout rates netNMF-sc had lower values of RMSE_0_. For example, at a dropout rate of 0.7, netNMF-sc had an average RMSE_0_ of 4.8 compared to 7.4 for the next best performing method, NMF (Figure 2(e)). Similar results were observed on data simulated using the *double exponential dropout model*. At a dropout rate of 0.7, netNMF-sc had an average RMSE_0_ of 8.3, slightly above MAGIC (average RMSE_0_ of 7.9) but substantially better than NMF (average RMSE_0_ of 15.9) and scImpute (average RMSE_0_ of 18.3) (Figure S6(e)). When we compute the RMSE between all entries of the transcript count matrix, scImpute outperforms other methods at low dropout rates (*<* 0.25) because scImpute does not attempt to impute non-zero entries. However, at dropout rates above 0.6, netNMF-sc has the lowest RMSE (Figure S4).

We investigated the contribution of the input network to the performance of netNMF-sc, and found that the addition of up to 70% random edges did not have a large effect on the performance (Figure S5).

### Evaluation on cell clustering

We compared netNMF-sc and other scRNA-seq methods in their ability to cluster cells into meaningful cell types using two scRNA-seq datasets. In both datasets, we normalized the count matrices following Zheng et al. [2017] to reduce the effect of differences in the total number of transcripts sequenced in each cell (library size) (See Supplemental section S1). We also applied a log-transformation (log_2_(**X** + 1)) to reduce the effect of outliers. We will refer to these datasets after normalization and log-transformation as the *raw data*.

The first data set contains 182 mouse embryonic stem cells (mESCs) that were flow sorted into one of three cell-cycle phases: G, S, and G2/M and sequenced using the Fluidigm C1 platform combined with Illumina sequencing [Buettner et al., 2015]. The raw data contains 9,571 genes and a dropout rate of 0.41. We computed cell clusters for each method as described above with two changes: (1) For netNMF-sc we used a network from the ESCAPE database [Xu et al., 2014] containing 153,920 protein-mRNA regulatory interactions from mESCs, with interaction weights of −1 for positive correlations and 1 for negative correlations. (2) We used *k* = 3 in *k*-means clustering to match the number of cell-cycle phases. We found that netNMF-sc outperformed other methods at clustering cells into the three cell cycle stages with an average adjusted Rand index (ARI) of 0.78 compared to 0.46 for MAGIC and 0.30 for scImpute (Figure 3, Figure S7). Note that while MAGIC did not perform as well as netNMF-sc in clustering the cells into distinct cell-cycle phases, it did identify a trajectory between the phases of the cell cycle, which may be biologically meaningful. However, notably MAGIC also identified a trajectory between clusters in the simulated data above although no trajectory was present (Figure 2(c)).

**Figure 3:**
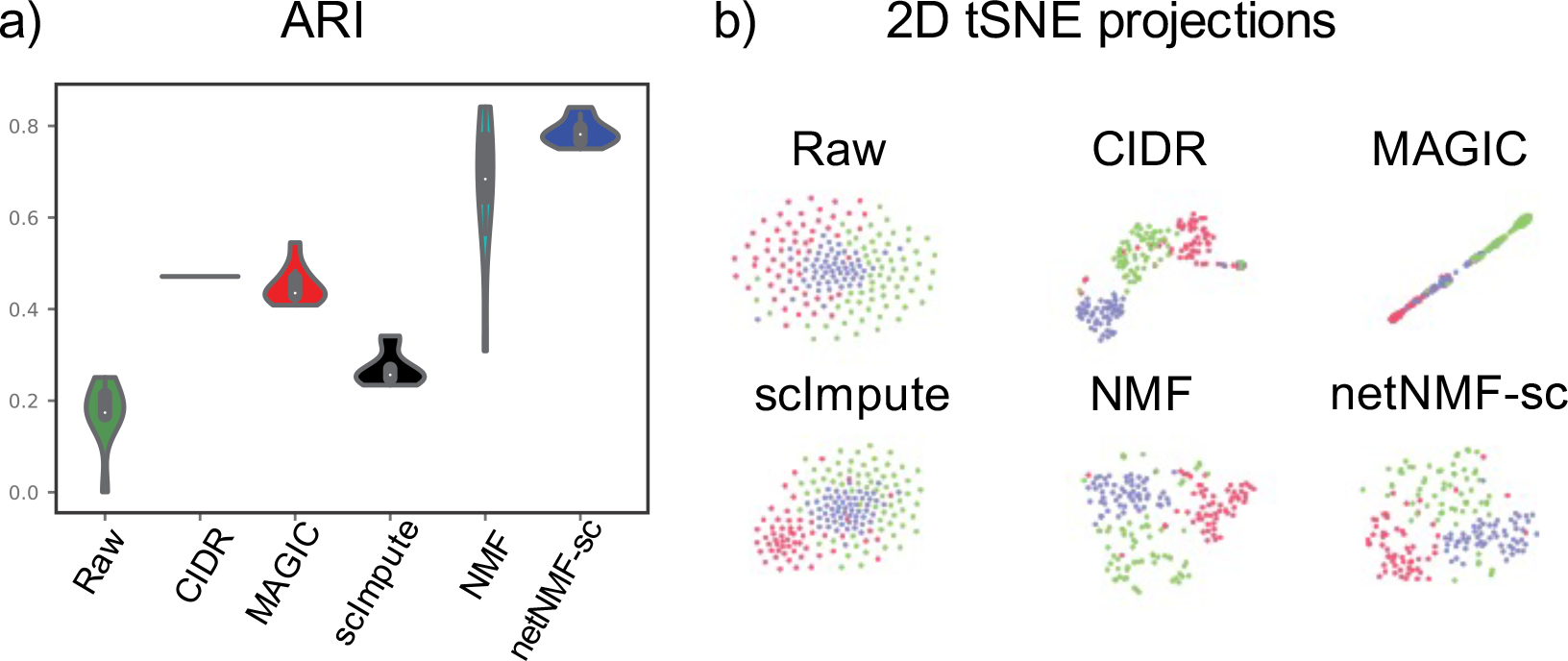
Clustering results on mouse embryonic stem cell (mESC) dataset from Buettner et al. [2015]. (a) The adjusted rand index (ARI) comparing cell cycle stages to cell clusters obtained from CIDR, from *k*-means clustering on the raw data, and from *k*-means clustering on imputed data from MAGIC, scImpute, NMF, and netNMF-sc. Each method was run 10 times, resulting in a distribution of ARI scores for all methods except for CIDR which is deterministic. (b) 2D *t*-SNE projections of imputed scRNA-seq data from each method.

To quantify the contribution of the network information to the performance of netNMF-sc, we ran netNMF-sc with three additional networks: a generic gene-gene co-expression network from Coexpedia [Yang et al., 2016], a *K*-nearest neighbors network (KNN), and a random network with the same degree distribution as the ESCAPE network. The *K*-nearest neighbors network was constructed by placing an edge between the ten nearest neighbors of each gene in the input data matrix **X**, based on Euclidean distance. We found that the ESCAPE co-expression network gave the best performance, with average ARI of 0.86 compared to 0.76 for Coexpedia, 0.73 for KNN, and 0.67 for the random network (Figure S7(f)). This result is not surprising as the ESCAPE network was constructed using only mESCs, while the Coexpedia network was constructed using cells from many different cell types. This demonstrates the benefit of prior knowledge that is matched to the tissue and cell types in the scRNA-seq data. Notably, all of the networks – including the random network – gave better performance than other methods suggesting that some of the advantage of netNMF-sc may be due to enforcing sparsity on **W**.

As a second comparison, we benchmarked netNMF-sc and other methods on a larger scRNA-seq dataset of 2022 brain cells from an E18 mouse sequenced using 10X Genomics scRNA-seq platform (https://support.10xgenomics.com/single-cell-gene-expression/datasets/2.1.0/neurons_2000). This dataset contains 13,509 genes with ≥ 10 transcript counts and a dropout rate of 0.84. We compared the cell clusters computed by each method with the 16 brain cell types reported in a separate 10X scRNA-seq dataset of 1.3 million cells from the forebrains of two different E18 mice that was analyzed using bigSCale [Iacono et al., 2018]. We ran each method as described above, with two changes: (1) for netNMF-sc we used a gene-gene co-expression network from [McKenzie et al., 2018] containing 157,306 gene-gene correlations across brain cell types (astrocytes, neurons, endothelial cells, microglia, and oligodendrodytes); (2) we used *k* = 16 in *k*-means clustering to match the number of brain cell types reported in bigSCale. We matched the cell clusters output by each method to the 16 cell types reported in bigSCale as follows. For each cluster, we computed the overlap between the top 200 over-expressed genes in each cluster (calculated as the log-fold change between cells in and out of the cluster) and the published marker genes for each of the 16 cell types. We then matched each cluster to the cell type with the lowest *p*-value of overlap (Fisher’s exact test). We marked a cluster as unclassified if the cluster was not enriched for any cell type with Bonferroni-corrected *p <* 0.05.

We found that the number of unclassified cells varied substantially across the methods. Clusters computed from the raw data, from MAGIC, and from NMF had 10%, 26%, and 1% of cells unclassified, respectively (Figure 4). In contrast, scImpute and netNMF-sc had no unclassified cells. While the true class assignment for each cell is unknown, we note that both scRNA-seq datasets were generated from the forebrains of E18 mice, and thus we expect that the proportions of each cell type should be similar across both datasets. We found that the proportions of each cell type identified by netNMF-sc were the closest (many within 2%) to the proportions reported by Iacono et al. [2018] (Figure 4f). In both cases, the cell type with the largest proportion is glutamatergic neurons, followed by interneurons and then radial glia and postmitotic neuroblasts. Other cell types, such as dividing GABAergic progenitors and Cajal-Retizus neurons, were found in smaller proportions. In contrast, MAGIC finds a large population (13%) of Cajal-Retizus neurons, while scImpute finds a large population (18%) of dividing GABAergic progenitor cells – both proportions more than 3-fold greater than in bigSCale or netNMF-sc. Clusters computed from the raw data and from NMF also differed substantially from the proportions reported in bigSCale; for example the proportion of post-mitotic neuroblasts was 0% in raw data, 20% in NMF, but 10% in bigSCale.

**Figure 4:**
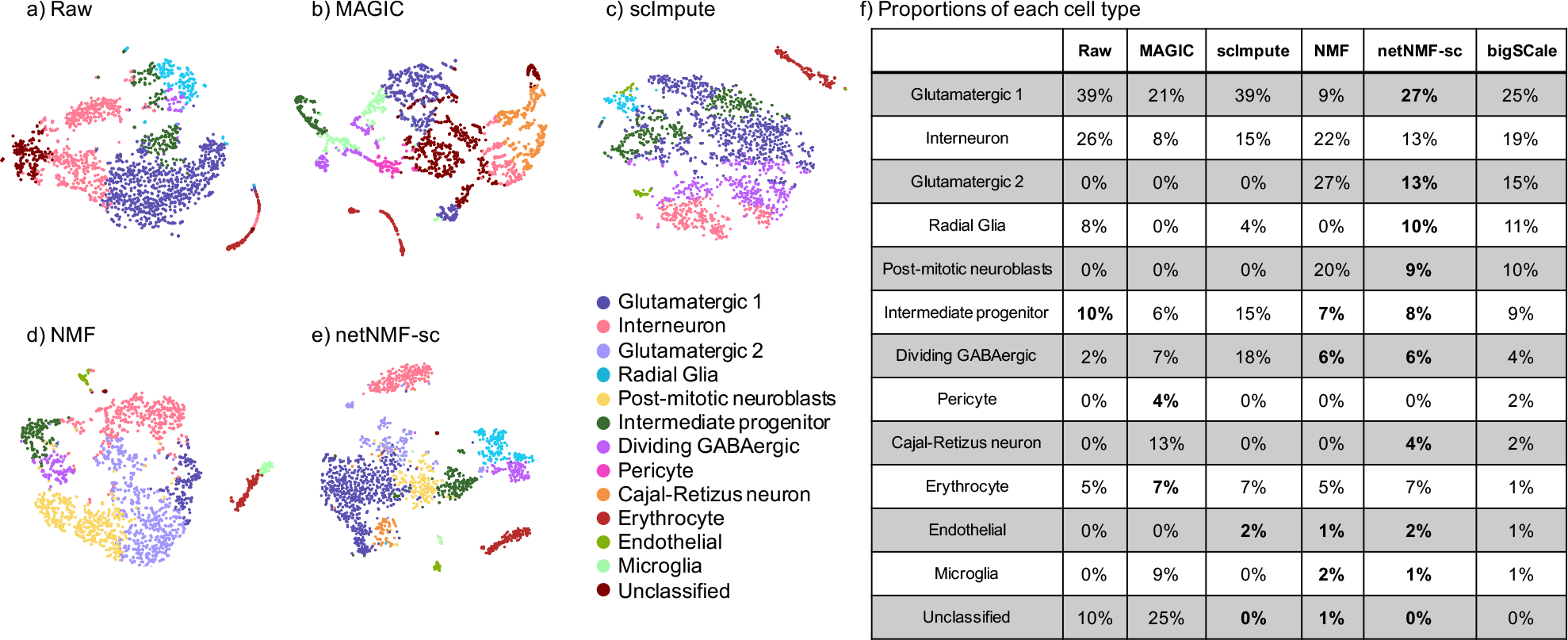
(a-e) *t*-SNE projections of raw and imputed scRNA-seq data from 2022 brain cells from an E18 mouse. Colors indicate cell types as derived in bigSCale analysis of 1.3 million E18 mouse brain cells [Iacono et al., 2018]. (f) Proportions of each cell type predicted by each method with p-values in parentheses. Entries in bold are within 2% of the proportions from bigSCale.

To further investigate the differences between clusters identified by netNMF-sc and other methods we examined the smallest cell cluster identified by netNMF-sc, containing only 14 cells. These cells were enriched (*p <* 1.3 × 10^*−*61^) for microglia marker genes reported by bigSCale, including well-studied marker genes such as *Csf1r*, *Olfml3*, and *P2ry12* [Wes et al., 2016] and represented 0.7% of the sequenced cell population, closely matching the proportion of microglia reported by bigScale (1%). NMF and MAGIC also identified clusters of microglia cells, but the differentially expressed genes in these clusters were less enriched for microglia marker genes (*p <* 4.1 × 10^*−*13^ and *p <* 5.5 × 10^*−*3^, respectively). The NMF cluster contained 65 cells but did not include any of the 14 cells classified as microglia by netNMF-sc, and was equally enriched for erythrocyte marker genes (*p <* 3.2 × 10^*−*11^). The MAGIC cluster contained 174 cells, a much larger proportion (9%) of the cell population than the 1% reported by bigSCale. This cluster included the 14 microglia identified by netNMF-sc but also 160 other cells. However, the additional 160 cells present in the cluster were not enriched for microglia marker genes (*p >* 1.2 × 10^*−*1^) but were enriched for glutamatergic marker genes (*p <* 1.5 × 10^*−*2^). This suggests that MAGIC erroneously grouped together different types of cells.

We next investigated the genes that weremost differentially expressed between the 14 microglia identified by netNMF-sc and the other 2008 cells. We found 436 genes to be significantly differentially expressed (FDR ≤ 0.01). All 50 microglia marker genes from bigScale were included in this set, including the two most highly differentially expressed genes *Cc14* (fold change 12.5) and *C1qc* (fold change 8.7). Of the top 20 differentially expressed genes several were reported in other studies [Sousa et al., 2017] but not bigScale, such as *Hexb* (fold change 7.8) and *Lgmn* (fold change 5.8). Among the top 20 differentially expressed genes were also several potential novel marker genes, including *Cstdc5* (fold change 4.5) and *Stfa1* (fold change 4.2) which have not been reported by other studies. These results suggest that netNMF-sc significantly improves clustering of cells into biologically meaningful cell types from scRNA-seq data with high dropout – even when the cell type is represented by only a small number (*<* 20) cells – and facilitates the identification of potential novel marker genes.

### Recovering marker genes and gene-gene correlations

We next investigated how well each method recovers differentially expressed marker genes and gene-gene correlations from scRNA-seq data. First, we examined cell-cycle marker genes. We obtained a set of 67 *periodic marker genes* whose expression has been shown to vary over the cell cycle across multiple cell types Dominguez et al. [2016]. This set contains 16 genes with peak expression in G1/S phase and 51 genes with peak expression during G2/M phase. We expect to observe a significant number of these periodic genes amongst the top differentially expressed genes between G1/S phase and G2/M phase cells in the cell cycle dataset from Buettner et al. [2015]. We compared the ranked list of differentially expressed genes from data imputed by netNMF-sc to the ranked lists of differentially expressed genes from the raw data and data imputed NMF, MAGIC, scImpute. We found that periodic genes ranked very highly in netNMF-sc results (*p* ≤ 3.2 × 10^*−*11^, Wilcoxon rank sum), a significant improvement compared to their ranking in the raw data (*p* ≤ 4.5 × 10^*−*3^, Wilcoxon rank sum) (Figure 5(a)). In contrast, imputing the data with NMF, MAGIC, and scImpute resulted in a lower ranking of the periodic genes (*p* ≥ 0.05, Wilcoxon rank sum). Additionally, we found that in data imputed by MAGIC, some periodic genes had expression patterns that were out of phase with the cell cycle. For example, *Exo1*, which peaks in G1/S phase, had significantly lower expression in G1/S phase cells compared to G2/M phase cells (*p* ≤ 2.2 × 10^*−*16^, Wilcoxon rank sum) in MAGIC imputed data (Figure 5(b)). In contrast, *Exo1* is not differentially expressed in the raw data (*p ≤* 0.17, Wilcoxon rank sum), but the peak in expression during G1/S phase is observed in the results from netNMF-sc (*p* ≤ 6.7 × 10^*−*12^, Wilcoxon rank sum) (Figure 5(b)).

**Figure 5:**
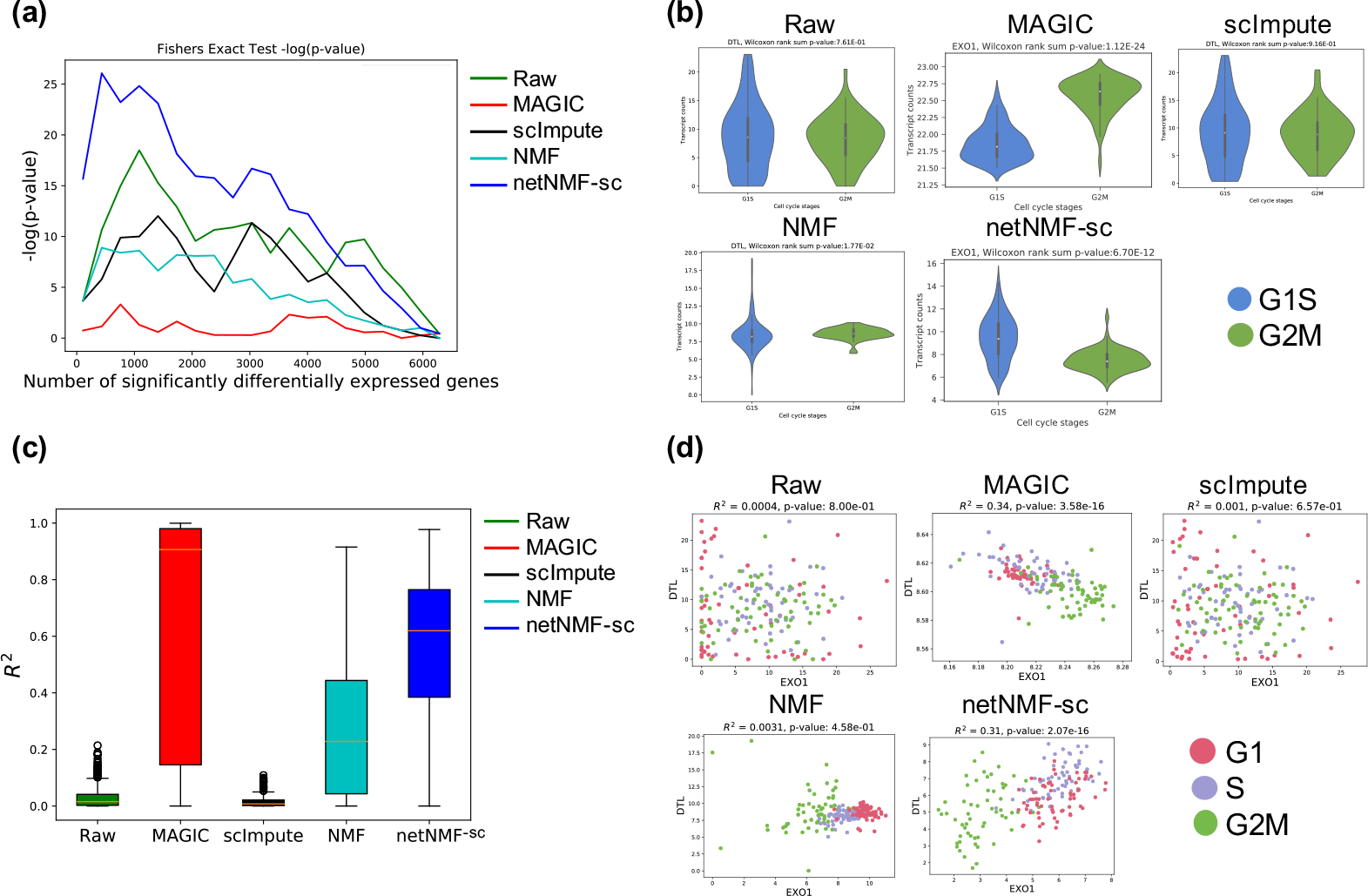
Comparison of differential expression of marker genes and gene-gene correlations in raw data from Buettner et al. [2015] and data imputed using netNMF-sc, NMF, scImpute, and MAGIC. (a) Overlap between differentially expressed genes and periodic genes (log *p*-values from Fisher’s exact test). (b) Expression of the G1/S phase marker gene *Exo1* in cells labeled as G1/S (blue) and cells labeled as G2/M (green) in data imputed by each method. In netNMF-sc inputed data, *Exo1* is overexpressed in G1/S cells compared to G2/M cells (*p* = 6.7 × 10^*−*12^), as expected. In contrast, in data imputed by MAGIC, *Exo1* is *underexpressed* in G1/S cells compared to G2/M cells (*p* = 1.12 × 10^*−*24^), and shows no significant difference in expression in raw and scImpute data. (c) Distribution of *R*^2^ correlation coefficients between pairs of periodic genes in cell cycle data. (d) Scatter plot of two G1/S phase genes: *Dtl* and *Exo1*. These genes are positively correlated in data imputed by netNMF-sc (*p* = 1.2 *×* 10^*−*19^), negatively correlated in data imputed by MAGIC (*p* = 3.6 *×* 10^*−*16^), and uncorrelated in other methods.

We also investigated whether each method could recover gene-gene correlations between periodic marker genes in the cell cycle data. We expect pairs of periodic genes whose expression peaks during the same phase of the cell cycle to be positively correlated and pairs of genes that peak during different phases to be negatively correlated. Across all 2211 possible pairs of periodic marker genes, we found that the mean *R*^2^ was 0.54 for netNMF-sc, compared to 0.73 for MAGIC, 0.29 for NMF, 0.02 for scImpute and 0.03 for raw data (Figure 5(c)). Setting a stringent cutoff for significant correlation (*R*^2^ ≥ 0.8*, p* ≤ 2.2 × 10^*−*16^), we found that 15% of periodic gene pairs were significantly correlated in data imputed by netNMF-sc compared to 68% in data imputed by MAGIC, 0.8% in data imputed by NMF, and nearly 0% in data imputed by scImpute. While the higher percentage of correlated gene pairs in MAGIC seems to be an advantage, it is important to recall that the MAGIC-imputed data contained marker genes, such as *Exo1*, whose expression signature was the *opposite* of expected. Such cases can result in incorrect correlations between pairs of marker genes. For example marker genes *Exo1* and *Dtl* both peak during G1/S phase and are expected to be positively correlated. However, MAGIC found significant negative correlation (*R* = −0.58*,p* = 3.6 × 10^*−*16^) between these two genes. In contrast, netNMF-sc recovers the positive correlation (*R* = 0.57*,p* = 2.1 × 10^*−*16^), while scImpute (*R* = 0.03*,p* = 0.66) and NMF (*R* = 0.045*,p* = 0.46) do not (Figure 5(d)).

Overall, we found that in the data imputed by MAGIC 19% of the significant gene pairs had correlations that were in the *opposite* direction than expected, i.e., genes that peaked during the same phase were negatively correlated and genes which peaked during different phases were positively correlated. In contrast, in the data imputed by netNMF-sc only 1% of the significant gene pairs had correlations in the opposite direction than expected (Figure 6(a)). These surprising results from MAGIC may be explained by the fact that MAGIC introduces a large number significant gene-gene correlation during imputation, many of which may be spurious, as was previously reported by Huang et al. [2018]. In fact, the majority (78%) of the gene-gene correlations in the correlation matrix generated from data imputed by MAGIC were significant (*R*^2^ ≥ 0.8*, p* ≤ 2.2 × 10^*−*16^), compared to only 0.2% in the correlation matrix generated from data imputed by netNMF-sc and 0.005% in the correlation matrix generated from the raw data (Figure 6(a)).

**Figure 6:**
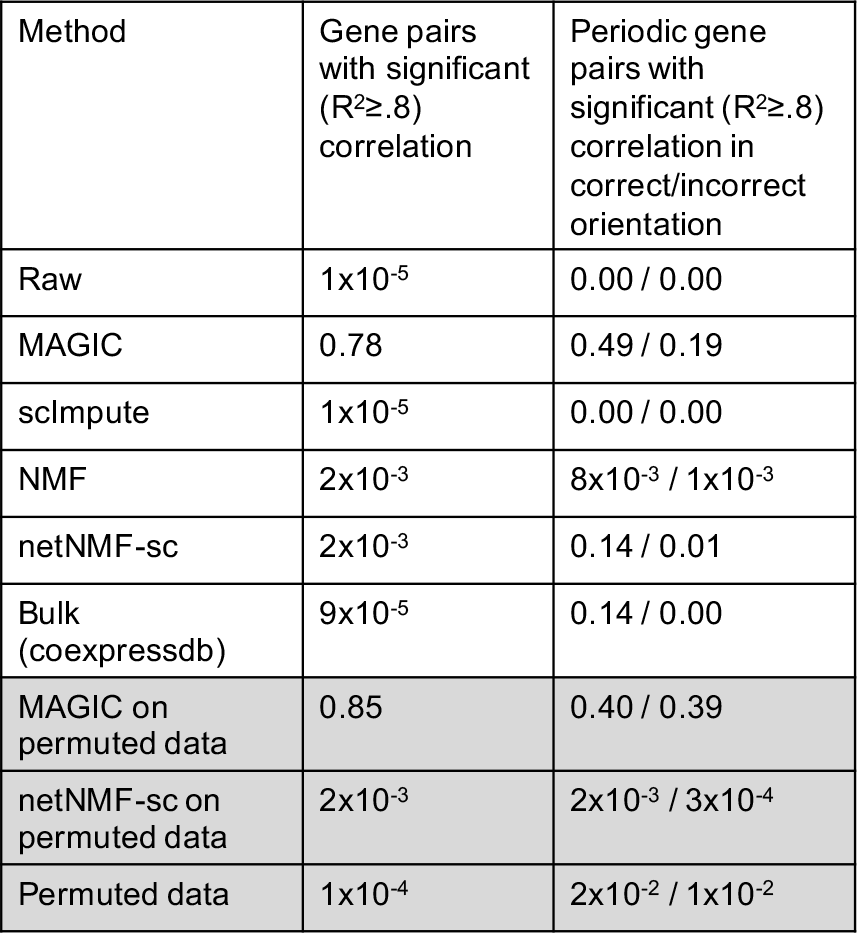
Fraction of all gene pairs and periodic gene pairs (defined by Dominguez et al. [2016]) with significant correlations (*R*^2^ ≥ 0.8, *p* ≤ 2.2 × 10^*−*16^ Student’s T-test) in the cell cycle dataset [Buettner et al., 2015]. *Correct* orientation means that a pair of genes with peak expression in the same stage of the cell cycle have positive correlation, and a pair of genes with peak expression in different stages of the cell cycle have negative correlation. Grey rows denote correlations on permuted data.

To examine whether these correlations identified by MAGIC and netNMF-sc represented real biological signal, we ran both methods on permuted data where the transcript counts of genes were permuted independently in each cell. We found that 85% of the gene pairs were significantly correlated (*R*^2^ ≥ 0.8*, p* ≤ 2.2 × 10^*−*16^) on MAGIC imputed data compared to only 0.2% of gene pairs on netNMF-sc imputed data (Figure 6). This observation suggests that many of the significant gene-gene correlations found in the MAGIC imputed cell cycle data may be spurious. Further investigation on simulated data suggests that such spurious correlations may be a consequence of the small number of cells: we found that MAGIC imputed data had many correlations in transcript count matrices with ≈ 200 cells but fewer correlations in imputed data with many (≈ 1000) cells (Figure S9). We also observed the number of gene-gene correlations found by MAGIC on permuted data increased rapidly with the diffusion parameter *t* before reaching a plateau, consistent with the diffusion process employed in MAGIC (Figure S8(b)). In contrast, the number of gene-gene correlations found by netNMF-sc on permuted data decreased as the number of latent dimensions *d* increased.

We performed a second analysis of differentially expressed marker genes and gene-gene correlations in scRNA-seq data from the MAGIC publication [Van Dijk et al., 2018] containing 7415 human transformed mammary epithelial cells (HMLEs) which were induced to undergo epithelial to mesenchymal transition (EMT) and then sequenced using the inDrops platform, a microfluidics based scRNA-seq technology [Klein et al., 2015]. We assessed how well each method recovered differential expression of 16 canonical EMT marker genes from Gibbons and Creighton [2018] (3 genes with high expression in epithelial (E) cells and 13 genes with high expression in mesenchymal (M) cells). We found that the EMT marker genes ranked very highly in netNMF-sc results (*p* ≤ 1.4 × 10^*−*5^, Wilcoxon rank sum), a significant improvement compared to their ranking in the raw data (*p* ≤ 3.1 × 10^*−*3^, Wilcoxon rank sum) (Figure S11(a)). In contrast, the next best performing method MAGIC had a smaller improvement in the ranking of EMT marker genes compared to the raw data (*p* ≤ 1.1 × 10^*−*4^, Wilcoxon rank sum). We observed that in data imputed by MAGIC, the E marker gene *TJP1* had higher average expression in M cells than E cells (*p* = 1.5 × 10^*−*33^) (Figure S11(b)). This resulted in *TJP1* being negatively correlated (*R* = −0.57*,p* = 3.4 × 10^*−*50^) with another E marker gene, *DSP* in the MAGIC imputed data; in contrast, these E marker genes showed positive correlation (*R* = 0.66*,p* = 6.4 × 10^*−*78^) in the netNMF-sc imputed data, correlation that was not apparent in the raw data (Figure S11(c)). We also investigated whether netNMF-sc could recover gene-gene correlations between EMT marker genes in E and M cells. We expect that pairs of E or M genes would exhibit positive correlation, while pairs containing one E and one M gene would exhibit negative correlations. In data imputed by netNMF-sc, 12% of the EMT gene pairs were significantly correlated (*R*^2^ ≥ 0.8*,p* ≤ 2.2 × 10^*−*16^), with all gene pairs correlated in the expected orientation (Figure S12). In data imputed by MAGIC, 23% of EMT gene pairs were significantly correlated, but 5% were correlated in the *opposite* direction than expected (Figure S11(d)). Full details of this analysis are in the Supplement.

## Discussion

We present netNMF-sc, a novel method for performing dimensionality reduction and imputation of scRNA-seq data in the presence of high (*>* 60%) dropout rates. These high dropout rates are common in droplet-based sequencing technologies, such as 10X, which is becoming the dominant technology for scRNA-seq. netNMF-sc leverages prior knowledge in the form of a gene-gene interaction network, and particularly a gene-gene coexpression network. Such networks are readily available for many tissue types, having been constructed from bulk RNA-sequencing data, or from other experimental approaches. To our knowledge, the only other other method that uses network information to perform dimensionality reduction on scRNA-seq data is Lin et al. [2017a]. However, this method assumes that there is no dropout in the data and its performance with high dropout rates is unknown. Moreover, this method uses a neural network that is trained on a specific protein-protein interaction (PPI) network, while netNMF-sc can use any gene-gene interaction network. Another method netSmooth [Ronen and Akalin, 2018] – published during the preparation of this manuscript – uses network information to smooth noisy scRNA-seq matrices but does not perform dimensionality reduction.

We demonstrate that netNMF-sc outperforms state-of-the-art methods in clustering cells in both simulated and real scRNA-seq data. In addition, netNMF-sc is better able to distinguish cells in different stages of the cell cycle and to classify mouse embryonic brain cells into distinct cell types whose proportions mirror the cellular diversity reported in a study with substantially greater number of cells. Similar to MAGIC, netNMF-sc imputes values for every entry in the input matrix, instead of imputing values only for zero entries as is done by scImpute. Imputation of all values can improve clustering performance and better recover biologically meaningful gene-gene correlations because transcript counts in scRNA-seq data are reduced for all genes. A potential downside of imputation is “oversmoothing” of the data resulting in the introduction of artificial gene-gene correlations. On multiple datasets, we show that netNMF-sc yields more biologically meaningful gene-gene correlations than other methods.

There are multiple directions for future improvement of netNMF-sc. First, netNMF-sc relies on existing gene-gene networks. While we have demonstrated that generic gene-gene co-expression networks [Yang et al., 2016] can improve clustering performance of human and mouse scRNA-seq data, netNMF-sc may not offer substantial improvements over existing methods on tissues or organisms where high-quality gene-gene networks are not available. In the future, other prior knowledge could be incorporated into netNMF-sc, such as cell-cell correlations, which might be obtained from underlying knowledge of cell types or from spatial or temporal information. Second, netNMF-sc relies on the parameter *λ* to weight the contribution of the network. Here we selected *λ* = 10 based on cross-validation on simulated data of different sizes, but a more rigorous procedure for selection of *λ* based on the input data and network may offer further improvement. Third, we have not fully explored how to select an appropriate dimension to use in netNMF-sc, and the choice of dimension may affect the quality of the imputation and clustering. Finally, there remains the issue of whether one should identify discrete cell clusters or continuous trajectories in scRNA-seq data. Here we focused on clustering cells in the the low-dimensional space obtained from netNMF-sc. An interesting future direction is to investigate how to leverage prior knowledge in trajectory inference from scRNA-seq data.

## Methods

### netNMF-sc algorithm

netNMF-sc uses graph-regularized NMF [Cai et al., 2008], which solves the following optimization problem:

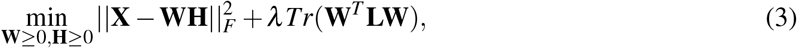

for a positive real constant *λ*, where **L** is the Laplacian matrix of the gene-gene interaction network, and *Tr*(⋅) indicates the trace of the matrix. The regularization term *Tr*(**W**^*T*^**LW**) encourages pairs of genes to be close in the matrix **W** when they are connected in the network. Graph-regularized NMF has previously been used in bioinformatics to analyze somatic mutations in cancer [Hofree et al., 2013].

Since we expect **X** to contain many zero entries that do not represent zero levels of that transcript but rather the effect of dropout, we formulate netNMF-sc such that a non-zero entry in *a*_*ij*_ of **WH** is not penalized when *x*_*ij*_ of **X** is equal to 0. We achieve this by using a binary matrix **M** to mask out zero entries. **M** has the same dimensions as **X** with entries

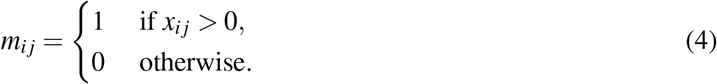

Incorporating the mask, the final formulation of netNMF-sc is

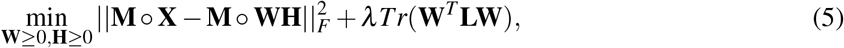

∘ indicates element-wise multiplication (or Schur product of matrices).

This formulation is similar, but not equivalent to a zero-inflated Poisson formulation of the counts [Lambert, 1992]. The zero entry mask is not implemented in many commonly used NMF methods [Pedregosa et al., 2011, MATLAB, 2018], but has a profound effect on improving clustering performance and imputation accuracy at high dropout rates (Figures S2(a-b)).

We derive the graph Laplacian **L** for the gene-gene interaction network as follows. Let **S** = [*s*_*ij*_] ∈ ℝ^*m×m*^ denote a gene similarity matrix whose entry *s*_*ij*_ is the weight of an interaction between genes *i* and *j*. A positive weight *s*_*ij*_ indicates a positive correlation between genes *i* and *j*, while a negative weight indicates negative correlation. We use the signed graph Laplacian **L** = **D** *−* **S**, where **D** = Diag(*|***S***|*1) is the degree matrix and *|***S***|* is the entry-wise absolute value of **S**. The signed Laplacian is symmetric and positive semidefinite, like the Laplacian [Kunegis et al., 2010, Gong et al., 2014]. Performing Laplacian embedding using the signed version of the graph Laplacian produces an embedding where positive edges between a pair of genes correspond to high similarity and negative edges correspond to low similarity [Kunegis et al., 2010].

We implemented netNMF-sc using the TensorFlow Python library [Girija, 2016] and tested the performance of netNMF-sc with four different optimizers: Adam, momentum, gradient descent, and Adagrad. We found Adam to perform the best at recovering clusters embedded in the data as well as reducing error between the imputed data and the original data prior to dropout (Figures S2(c-d)). Adam (Adaptive Moment Estimation) uses the first and second moments of gradient of the cost function to adapt the learning rate for each parameter [Kingma and Ba, 2014]. This allows Adam to perform well on noisy data as well as sparse matrices [Kingma and Ba, 2014].

netNMF-sc runs in 1.8 minutes on one 2.60GHz Intel Xeon CPU (or in 41 seconds on one NVidia Tesla P100 GPU) on a dataset with 10,000 genes and 1000 cells. In comparison, scImpute required 3.6 minutes on the same CPU and dataset. (Figure S3).

### Generation of simulated scRNA-seq data

We used the simulator SPLATTER [Zappia et al., 2017] to generate complete transcript count data, estimating the parameters of the model from mouse embryonic stem cell scRNA-seq data [Buettner et al., 2015] using the SplatEstimate command. We modified SPLATTER to introduce correlations between genes that are differentially expressed in each cluster using a gene coexpression network from Yang et al. [2016]. See Supplemental Section S3 for further details.

After simulating transcript counts to obtain a complete count matrix **X**′, we generated dropout events using one of two models. The first is a *multinomial dropout model*, used previously to model dropout in scRNA-seq data Linderman et al. [2018], Zhu et al. [2018]. In this model, the observed counts in each cell are multinomial distributed, where the probability of observing a transcript from gene *i* in cell *j* is 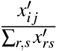 and the number of trials is the sum of all transcripts in the complete count matrix,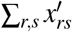, multiplied by the capture efficiency, ranging from 0 to 1. The resulting count matrix **X** contains dropout proportional to the capture efficiency. The second model is the *double exponential dropout model*, used previously in the scImpute [Li and Li, 2018] and BISCUIT [Azizi et al., 2017] publications. In this model, an entry *x*_*ij*_ of the count matrix is set to zero with probability 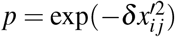, where *δ* is the dropout rate.

## Availability

netNMF-sc is available at github.com/raphael-group/netNMF-sc.

## Acknowledgements

This project has been made possible in part by grant numbers 2018-182608, 1005664, and 1005667 from the Chan Zuckerberg Initiative DAF, an advised fund of Silicon Valley Community Foundation. BD and BEE were also funded by NSF CAREER 1750729. BJR was also funded by NSF CAREER CCF-1053753.

